# Chromatin accessibility dynamics of myogenesis at single cell resolution

**DOI:** 10.1101/155473

**Authors:** Hannah A. Pliner, Jonathan Packer, José L. McFaline-Figueroa, Darren A. Cusanovich, Riza Daza, Sanjay Srivatsan, Xiaojie Qiu, Dana Jackson, Anna Minkina, Andrew Adey, Frank J. Steemers, Jay Shendure, Cole Trapnell

## Abstract

Over a million DNA regulatory elements have been cataloged in the human genome, but linking these elements to the genes that they regulate remains challenging. We introduce Cicero, a statistical method that connects regulatory elements to target genes using single cell chromatin accessibility data. We apply Cicero to investigate how thousands of dynamically accessible elements orchestrate gene regulation in differentiating myoblasts. Groups of co-accessible regulatory elements linked by Cicero meet criteria of “chromatin hubs”, in that they are physically proximal, interact with a common set of transcription factors, and undergo coordinated changes in histone marks that are predictive of gene expression. Pseudotemporal analysis revealed a subset of elements bound by MYOD in myoblasts that exhibit early opening, potentially serving as the initial sites of recruitment of chromatin remodeling and histone-modifying enzymes. The methodological framework described here constitutes a powerful new approach for elucidating the architecture, grammar and mechanisms of *cis-*regulation on a genome-wide basis.

## Introduction

Chromatin accessibility is a powerful marker of active regulatory DNA. In eukaryotes, chromatin accessibility at both gene promoters and distal regulatory elements, many of which are enhancers, delineates where DNA binding factors are bound in place of nucleosomes(Felsenfeld et al., 1996). Genome-wide analyses of chromatin accessibility as measured by DNaseI hypersensitivity have found that the repertoire of accessible regulatory elements in cell lines or tissues constitutes a highly specific molecular signature (Thurman et al., 2012). Furthermore, genome-wide association studies (GWAS) consistently show that a substantial proportion of genetic risk for common disease falls within regions exhibiting chromatin accessibility in disease-relevant tissues or cell types (Gusev et al., 2014; Maurano et al., 2012).

Despite the clear importance of chromatin accessibility, we lack a quantitative understanding of how changes in chromatin accessibility relate to changes in the expression of nearby genes. A first challenge is to generate a map that links distal regulatory elements with their target genes, as enhancers can operate at long distances and influence multiple genes. A second challenge is to understand the determinants of that map. What factors govern which gene(s) an enhancer will regulate, what is the degree of activation that an enhancer will confer, and what are the interactions and dependencies of an enhancer with respect to other regulatory elements acting on the same gene(s)?

Here, we describe Cicero, an algorithm that links distal regulatory elements to their target genes based on patterns of co-accessibility in single cell chromatin accessibility data. In contrast with previous approaches that rely on a large compendium of bulk chromatin accessibility data generated across many cell lines or tissues (Thurman et al., 2012) (see Budden et al., 2015) for a review of recent approaches), Cicero uses single cell chromatin accessibility data from only a single experiment. Cicero identifies functional regulatory elements and links them with target genes using unsupervised machine learning. The algorithm can therefore be applied to any cell type and organism for which a sequenced genome and single cell chromatin accessibility data are available. Because it accepts single cell data as input, Cicero can in principle work on complex mixtures of different cell types as are found in tissues.

We demonstrate Cicero’s capabilities through an analysis of skeletal myoblast differentiation, which remains one of the best characterized model systems for gene regulation in vertebrate development. Myoblast differentiation is orchestrated by a core set of transcription factors, including MYOD and the MEF2 proteins (Molkentin et al., 1995), which regulate the expression of thousands of genes as cells exit the cell cycle, align, and fuse to form myotubes. We identified over 300,000 accessible DNA elements in myoblasts, nearly 12,000 of which open or close during differentiation. When applied to single cell combinatorial indexing ATAC-seq (sciATAC-seq) data generated from thousands of differentiating myoblasts, Cicero linked the vast majority of dynamic sites to one or more target genes. From the resulting *cis*-regulatory map, we can predict changes in gene expression based on the chromatin accessibility dynamics of the linked distal elements.

Co-accessible sites linked by Cicero bear the hallmarks of “chromatin hubs”, *i.e.* a spatial configuration wherein distal DNA regulatory elements are looped into the vicinity of their target genes. Through comparison with chromosome conformation capture data, we demonstrate that accessible elements linked by Cicero are in physical proximity. We also observe coordinated deposition and removal of histone marks at linked sites, suggesting that histone-modifying enzymes recruited to one site modify histones at other sites in the chromatin hub. For example, histone acetylation near enhancers frequently co-occurred with acetylation at target promoters and downstream of transcription start sites (TSSs). Sequence motifs within dynamically accessible distal sites predicted changes in histone modifications and expression at linked genes. Our observations are consistent with a model in which recruitment of epigenetic modifiers to one more or sites in a chromatin hub propagates changes to all of them, facilitating coordinated regulation of target genes.

## Results

### The trajectories of chromatin accessibility and gene expression during myoblast differentiation are highly similar

We performed a differentiation time course on human skeletal muscle myoblasts (HSMM), harvesting cells at 0, 24, 48 and 72 hours after the switch from growth media to differentiation media (**Figure 1A**). Using an improved version of our sci-ATAC-seq protocol (Cusanovich et al., 2015), we profiled chromatin accessibility in a total of 5,060 cells, of which 3,053 were triaged as potentially corresponding to interstitial fibroblasts (see Methods; a similar proportion were triaged in single cell RNA-seq datasets generated in the same system (Qiu et al., 2017)). Upon pooling reads from the remaining 2,007 differentiating myoblasts and calling peaks using MACS 2 (Feng et al., 2012), we identified 326,829 accessible sites, covering 5.7% of the genome. Of these, 34% overlapped DNaseI hypersensitive sites (DHSs) identified in myoblasts or myotubes and 81% overlapped combined DNaseI hypersensitive sites from 125 cell lines by ENCODE (The ENCODE Project Consortium, 2012) (**Supplemental Figure 1A**). The overwhelming majority of sites detected by sci-ATAC-seq that were not identified by DNaseI mapping were detected in fewer than 100 cells (**Supplemental Figure 1B**). Each cell had reads overlapping with, on average, 2,842 promoter-proximal accessible sites and 7,534 distal accessible sites (**Supplemental Figure 1C**).

**Figure 1.**
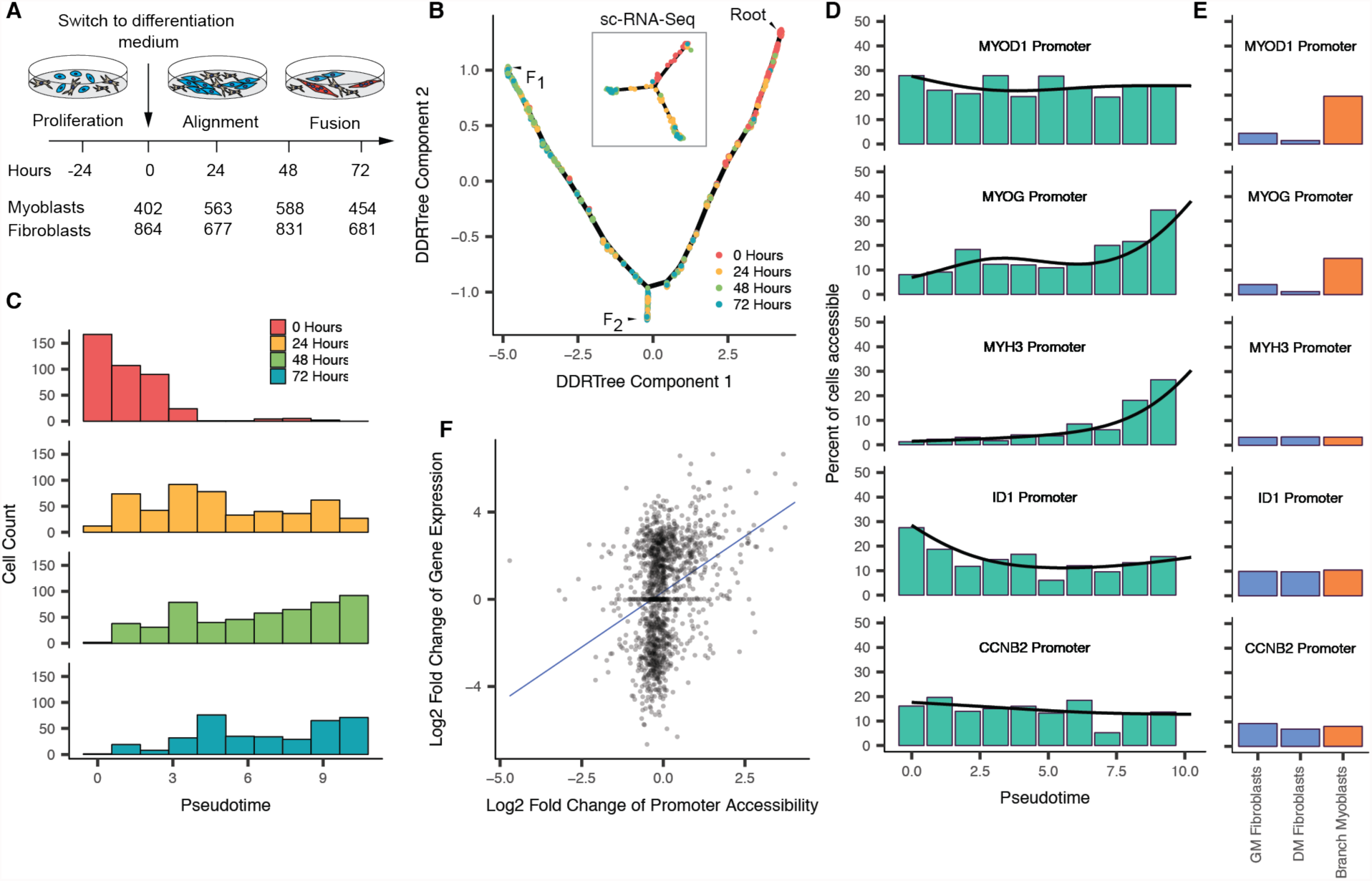
Differentiating myoblasts follow similar single cell chromatin accessibility and gene expression trajectories. **A**) Single cell chromatin accessibility profiles for human skeletal muscle myoblasts (HSMM) were constructed with sci-ATAC-seq. Contaminating interstitial fibroblasts (common in HSMM cultures) were removed informatically prior to further analysis. **B**) The single cell trajectory inferred from 2,007 myoblast sci-ATAC-seq profiles by Monocle (see Methods). In subsequent panels and throughout the paper, we exclude cells on the branch to outcome F_2_ unless otherwise indicated, leaving 1,797 cells on the root and F_1_ branches. Inset shows the sc-RNA-seq trajectory reported for HSMMs (reproduced from Figure 2 of Qiu et al. (2017)). **C**) Distribution of cells in chromatin accessibility pseudotime from the root to trajectory outcome F_1_. **D**) Percent of differentiating cells whose promoters for selected genes are accessible across pseudotime. Black line indicates the pseudotime-dependent average from a smoothed binomial regression. **E**) Percent of cells whose promoters for selected genes in D are accessible in fibroblasts collected in growth medium (GM) or differentiation medium (DM), as well as myoblasts localized to the branch to F_2_. **F**) Change in promoter accessibility versus change in gene expression. Log2 fold changes were calculated by dividing the pseudotime-dependent average from a smoothed negative binomial regression at the 90th percentile of pseudotime by the average at the 10th percentile of pseudotime. A pseudocount of 0.1 was added to each prior to computing the fold changes.

We next sought to characterize changes in chromatin accessibility as myoblasts differentiated, and to relate these accessibility changes to changes in gene expression. However, analyzing cell differentiation at single cell resolution is challenging in part because cells execute programs of gene regulation asynchronously. To overcome this problem, we recently developed a technique called “pseudotemporal reordering” (or “pseudotime”) that organizes individual cells using machine learning according to their progress through a biological process such as cell differentiation (Trapnell et al., 2014). Although our algorithm, Monocle, was designed to order single cells based on their transcriptomes (Qiu et al., 2017), we were able to adapt it to sci-ATAC-seq data with straightforward modifications (see Methods).

Monocle placed the cells along a trajectory with two outcomes (denoted F_1_ and F_2_) that was strikingly similar to the trajectory constructed from single cell transcriptome data (**Figure 1B**). Cells harvested from growth media fell almost exclusively near the beginning of the trajectory, while cells from later time points were distributed over its length (**Figure 1C**). Over the path to F_1_, promoters for well-known myogenic regulators and structural components of muscle opened (*i.e.* became more accessible), whereas the promoter of ID1, a well-characterized repressor of myoblast differentiation (Benezra et al., 1990), closed (**Figure 1D**). Similar to the single cell RNA-seq trajectory (Qiu et al., 2017), a substantial number of cells were positioned on a short branch leading to the alternative outcome F_2_. Although these potentially correspond to interstitial fibroblasts that we failed to filter, their levels of accessibility in the *MYOD1* promoter are similar to differentiating myoblasts. Nevertheless, the promoters of *MYOG*, *MYH3*, and other markers of fully differentiated myocytes did not significantly open in this branch (**Figure 1E**), consistent with these representing “reserve myoblasts” that did not fully differentiate (Friday and Pavlath, 2001; Yoshida et al., 1998).

The similar trajectories constructed by Monocle from single cell RNA-seq and sci-ATAC-seq data, and the close correspondence between the kinetics in expression and chromatin accessibility for key muscle genes, support the conclusion that Monocle’s pseudotime ordering is accurate. Globally, however, changes in promoter accessibility correlated poorly with changes in expression (**Figure 1F**). For example, accessibility for the promoter of CCNB2, which is transcriptionally downregulated in differentiating HSMMs, remained mostly constant over pseudotime (**Figure 1D**).

### Distal DNA elements are dynamically accessible during myoblast differentiation

Differential analysis revealed significant pseudotime-dependent changes in accessibility at 11,948 of 326,829 (3.7%) sites during myoblast differentiation (**Figure 2A**). Of these “dynamic” sites, only 736 (6.2%) were promoters (**Figure 2B**), of which 67 overlapped with 1,464 differentially expressed transcripts (FDR < 5%). Of the 51 of these promoters with stable accessibility and gene expression change (i.e. not transient change), 48 (94%) were directionally concordant (increase in accessibility and expression or decrease in both). 72% of the remaining 11,212 distal dynamically accessible sites were annotated as enhancers in myoblasts or myotubes by Segway (Libbrecht et al., 2016), as compared with only 32% of all accessible sites (**Figure 2B**).

**Figure 2.**
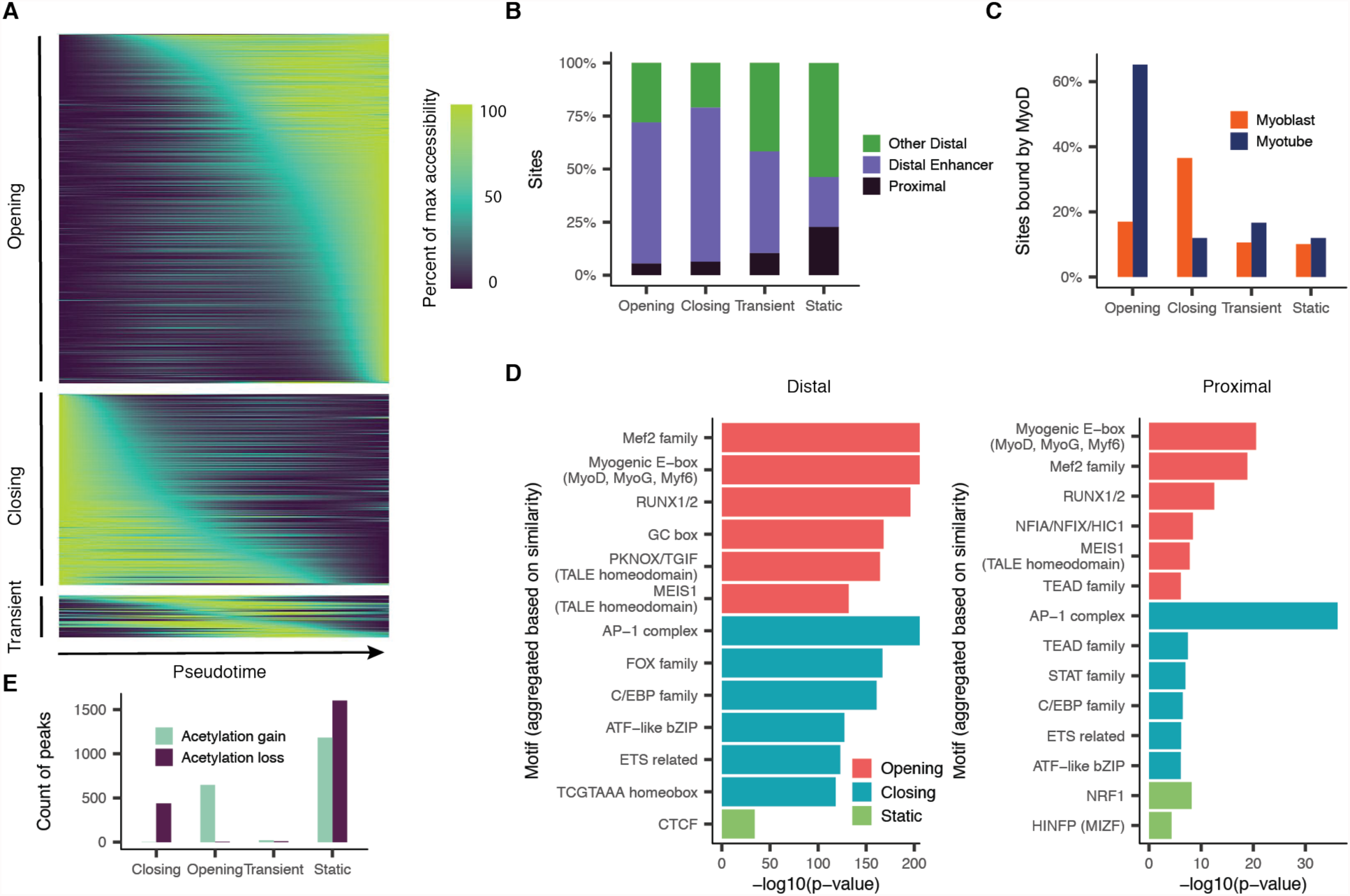
Thousands of DNA elements are dynamically accessible during myoblast differentiation. **A**) Smoothed pseudotime-dependent accessibility curves, generated by a negative binomial regression and scaled as a percent of the maximum accessibility of each site. Each row indicates a different DNA element. Sites are sorted by the pseudotime at which they first reach half their maximum accessibility. **B**) Proportions of dynamic and static sites by site type. Color indicates whether a site is promoter-proximal (defined as a peak overlapping the first 500 bp upstream of a TSS, see Methods), a distal enhancer (defined as peaks that are not promoter-proximal, and are annotated by Segway as enhancers in either myoblasts or myotubes), or other distal (defined as distal sites not annotated as enhancers by Segway). **C**) Percent of sites reported as bound by MyoD in either myoblasts or myotubes by Cao et al. (2010) **D**) Motif enrichments in accessible sites. P-values result from logistic regression models that use the presence or absence of a given motif in each site to predict whether the site has a given accessibility trend (opening/closing/static). Each motif is tested against each of the three accessibility trends separately. Plots show up to the top 6 Bonferroni-significant motifs by p-value. **E**) Counts of sites undergoing significant changes in H3K27 acetylation as measured by ChIP-Seq (Tang et al., 2015).

Using gene set enrichment analysis, we found that genes associated with contraction and other muscle-related functions were strongly enriched among genes with significantly opening promoter regions. In contrast, promoters for genes associated with the cell cycle, which are downregulated early in differentiation, were only marginally enriched among the differentially accessible sites (**Supplemental Figure 2A**). Like CCNB2, most markers of actively proliferating cells did not show significant changes in promoter accessibility. A potential explanation is that cyclins are downregulated by direct binding of Rb to E2F family proteins at their transactivation domains (Weinberg, 1995), reducing their expression without requiring nucleosome deposition at their promoters.

A broader analysis of transcription factor binding at dynamically accessible sites showed that most changes are concomitant with the redistribution of known myogenic regulators. Comparison to ChIP-seq data collected by Tapscott and colleagues(Cao et al., 2010) revealed that 65% of opening sites and 36% of closing sites are bound by MYOD in myotubes and myoblasts respectively (**Figure 2C**). In contrast, only 15% of static sites (those without significant changes in accessibility) were MYOD-bound in either myoblasts or myotubes. Dynamic distal elements and promoters were also strongly enriched for binding motifs for MYOD, MYOG, and MEF2 family members and other transcription factors with central regulatory roles in myogenesis (**Figure 2D**).

Many transcription factors, in addition to their role in nucleating the assembly of the pre-initiation complex, also function to recruit enzymes that mark histones near regulatory DNA elements with modifications associated with active expression. For example, MYOD recruits p300(Sartorelli et al., 1997), whose histone acetyltransferase activity is required for its role in activating gene expression (Dilworth et al., 2004; Puri et al., 1997; Sartorelli et al., 1997). A comparison with ENCODE data in myoblasts and myotubes showed overwhelming directional concordance between sites that were gaining or losing H3K27 acetylation (H3K27ac) vs. sites that were opening or closing in chromatin accessibility, respectively (**Figure 2E**). However, the vast majority of changes in histone marks during differentiation cannot be explained by changes in chromatin accessibility. For instance, the integrative chromatin state catalog constructed by Libbrecht et al. (2016) shows histone modification state switching among 63% of statically accessible sites (**Supplemental Figure 2C**). Thus, myoblast differentiation is characterized by the modulation of histone marks at hundreds-of-thousands of sites, only a minority of which can be explained by changes in chromatin accessibility.

### Cicero constructs genome-wide cis-regulatory models from sci-ATAC-seq data

We sought to exploit patterns of co-accessibility between distal regulatory elements and promoters to build a genome-wide cis-regulatory map from sci-ATAC-seq data. However, constructing a co-accessibility map is challenging for several reasons. First, the raw correlations in accessibility between sites are driven in part by technical factors such as read depth per cell. Second, we do not have sufficient observations to accurately estimate correlations between billions of pairs of sites. Finally, while the accessibility of regulatory elements might be correlated with their target promoters, pairs of very distant sites, including those on other chromosomes, will also be correlated, by virtue of being part of the same gene regulatory program. For example, one might expect most MYOD-bound sites to be correlated to some degree during myogenesis.

To address these challenges, we developed a new algorithm, Cicero, that subtracts technical and genomic distance effects while constructing a global cis-regulatory map from single cell chromatin accessibility profiles (**Figure 3A**). First, the algorithm groups cells by their state using unsupervised clustering or by position along a pseudotemporal trajectory. Second, it aggregates accessibility profiles for cells in these groups to produce counts that can be readily adjusted to subtract the effects of experimental batch, overall number of observed sites, and other technical variables. Third, it computes the correlations in adjusted accessibilities between all pairs of sites within 500 kilobases (kb) of one another. To calculate robust correlations, we use a statistical machine learning technique called the Graphical LASSO (Friedman et al., 2008), which estimates regularized correlation matrices. Cicero penalizes correlations between sites that are far apart in the genome more than those that are close together, preserving local patterns of co-accessibility at the expense of very long range ones. The output of Cicero is a set of all pairs of sites in the genome within 500 kb of each other, along with their co-accessibility scores.

**Figure 3.**
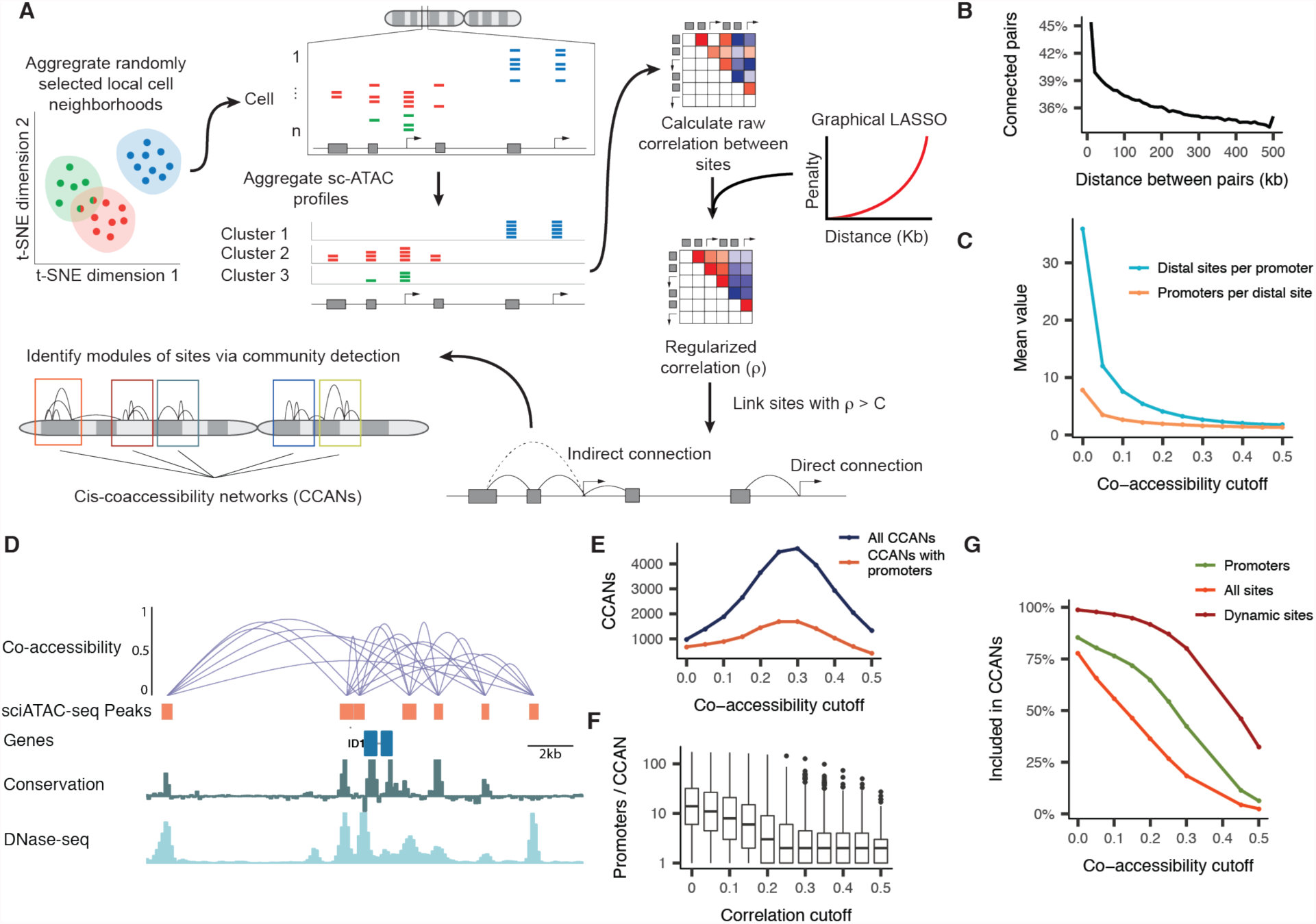
Cicero constructs cis-regulatory models genome-wide from sci-ATAC-seq data. **A**) An overview of the Cicero algorithm (see Methods for details) **B**) Fraction of pairs of sites with positive co-accessibility scores at varying distances in the linear genome. **C**) The mean number of distal sites per promoter and promoters per distal site at increasing co-accessibility score cutoffs. **D**) A summary of the Cicero co-accessibility network containing the *ID1* locus. Connections to elements outside the window shown are excluded for clarity. The height of connections indicates the magnitude of the Cicero co-accessibility score between the connected peaks. The conservation track shows the 100 vertebrate PhyloP conservation scores (phyloP100wayAll). The DNase-seq track shows the ENCODE signal track for HSMM. **E**) Number of CCANs with 3 or more co-accessible DNA elements identified with Louvain community detection. Prior to running Louvain, connections below the indicated Cicero co-accessibility score were excluded (see Methods for details). CCANs that include the promoter of at least 1 detectably expressed gene are shown as a separate series. **F**) Boxplots of the number of expressed gene promoters per CCAN at increasing co-accessibility score cutoffs. **G**) Percent of sites recruited into a CCAN at increasing co-accessibility score cutoff. Colors represent subsets of sites: green represent promoters for genes that are detectably expressed; orange and red represent sites that are accessible and differentially accessible across pseudotime, respectively.

We applied Cicero to generate a genome-wide cis-regulatory map from our sci-ATAC-seq data from 1,797 differentiating myoblasts. Cicero identified 6,424,549 pairs of sites with positive co-accessibility scores, including 1,680,416 pairs between a promoter and a distal element. (**Figure 3B**). The magnitude of a Cicero co-accessibility score indicates the strength of co-accessibility across cells. As the co-accessibility threshold is raised, promoters are connected to fewer, high-confidence regulatory elements. For example, at a co-accessibility score cutoff of 0.25, promoters were, on average, connected to 3 distal elements, and each distal element was linked to 1.7 promoters (**Figure 3C**). Cicero connected the promoter of *ID1*, a negative regulator of the myogenic program that is transcriptionally silenced during differentiation, to 4 conserved, distal regulatory elements within 10 kb (**Figure 3D**).

We noticed that co-accessible DNA elements tended to cluster together in the genome, so we post-processed Cicero’s co-accessibility output with a community detection algorithm to identify “cis-co-accessibility networks” (CCANs): modules of sites that are highly correlated with one another. Cicero allows a user to set a threshold co-accessibility that controls the size and granularity of CCANs (**Figure 3E-G**). At lower co-accessibility thresholds, sites were linked into only a few very large CCANs, while at high co-accessibility thresholds, most sites were isolated and most genes were disconnected from putative regulatory elements (**Figure 3E**). At a co-accessibility score threshold of 0.25, we identified 4,481 CCANs which incorporated 87,574 sites. Ultimately, we chose this threshold to maximize both the number of CCANs identified and the number of sites included in the CCANs. On average, each CCAN involved 575 kb of the linear genome, contained 19 sites, and included the promoters of 2.2 expressed genes (**Figure 3F**). Overall, 57 percent of expressed genes, 27% of all peaks, and 87% of differentially accessible peaks are included in Cicero’s CCANs (**Figure 3G**). CCANs included the promoters for 824 of the 1,464 genes differentially expressed during myoblast differentiation.

We hypothesized that CCANs linked by Cicero constitute “chromatin hubs”. Chromatin hubs, which are thought to involve looping interactions between distal regulatory elements and the genes they target, may act to coordinate the assembly of transcription complexes (de Laat and Grosveld, 2003; Tolhuis et al., 2002). To satisfy the definition of a chromatin hub, we expect that groups of genomic sites linked by Cicero should meet four criteria. First, they should exhibit greater physical proximity than expected based on their distance in the linear genome. Second, they should interact with a common set of protein complexes. Third, they should be epigenetically modified in concordant ways and at similar times. Finally, they should substantively contribute to regulating genes with promoters within the hub.

### Co-accessible DNA elements exhibit physical proximity

We asked whether co-accessible sites are closer together in the nucleus than unlinked sites at similar distances in the linear genome. Previous analyses of ENCODE cell lines reported that distal DHSs and promoters that display a correlation in their sensitivity to DNase I were closer in the nucleus according to chromosome conformation capture data than uncorrelated pairs of sites (Malin et al., 2013; Thurman et al., 2012). To test this in Cicero-based links, we used our updated sci-ATAC-seq protocol to generate chromatin profiles from 662 human lymphoblastoid cells (GM12878), for which high-resolution Hi-C data was recently generated (Rao et al., 2014). Pairs of sites linked by Cicero were in contact more frequently than unlinked sites at similar genomic distances (**Figure 4A**).

**Figure 4.**
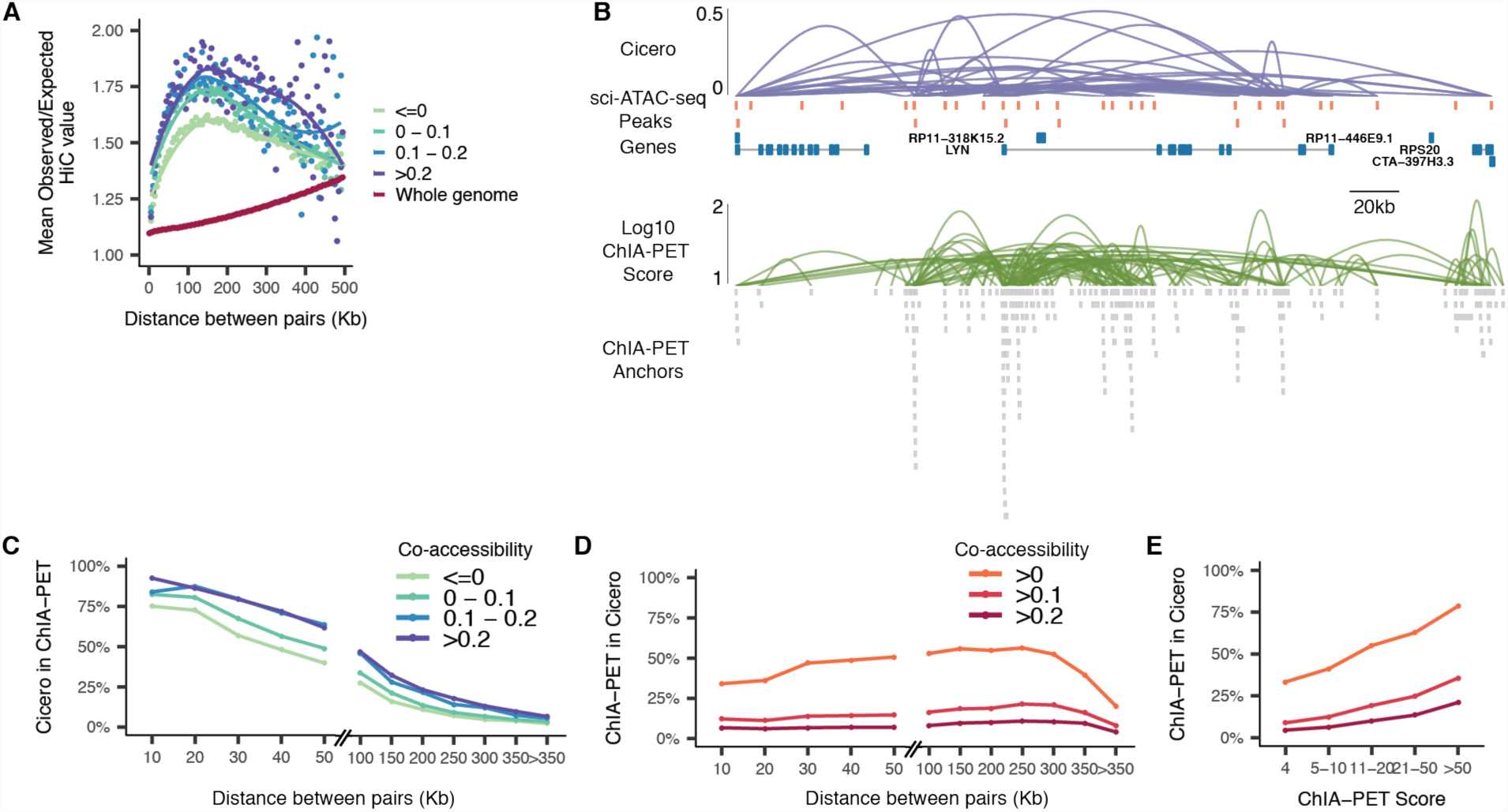
Co-accessible DNA elements linked by Cicero are physically proximal in the nucleus. **A**) Observed contact frequencies for pairs of sites linked by Cicero at varying distances in the linear genome, normalized to expected contact frequencies. Sites not connected by Cicero are shown for comparison. “Whole genome” indicates the observed:expected ratio for all pairs of genomic bins. **B**) Cicero connections for the LYN locus compared to RNA pol-II ChIA-PET data. **C**) CCAN connections detected in ChIA-PET contacts as a function of distance. Color indicates the magnitude of the Cicero co-accessibility score. **D**) ChIA-PET contacts found in Cicero connections as a function of distance. Colors in D and E indicate the magnitude of the Cicero co-accessibility scores included. **E**) ChIA-PET contacts detected in Cicero CCAN connections as a function of ChIA-PET score.

We also observed strong concordance between Cicero-based linkages and DNA elements in RNA pol II-mediated contacts captured via ChIA-PET (Tang et al., 2015). About half of DNA elements ligated via ChIA-PET (“anchors”) overlapped with accessible sites in our data, with greater overlap between anchors that were supported by multiple ChIA-PET reads and sites that were accessible in many cells (**Supplemental Figure 3**). For example, Cicero constructed a regulatory map surrounding the *LYN* locus that was strikingly similar to the pol II-mediated contact map produced by ChIA-PET (**Figure 4B**). Altogether, 26% (n = 37,823) of pairs of anchors that were linked by Cicero were also found to be in contact via ChIA-PET, with greater concordance for linkages with higher Cicero co-accessibility (**Figure 4C**). Although agreement between pol II ChIA-PET and Cicero linkages was substantially higher for closely located sites, the assays exhibited concordance even for sites separated by 100 kb or more. Furthermore, 46% (n = 16,066) of pairs of accessible sites found to be in contact via ChIA-PET were also linked by Cicero, with greater concordance for higher confidence ChIA-PET connections (**Figure 4D-E**). Taken together, these analyses support the view that sites that are co-accessible within single cells are in physical proximity in the nucleus, even when separated by tens to hundreds of kilobases in the linear genome.

### Co-accessible DNA elements carry pairs of motifs for interacting transcription factors

Concordance between pairs of sites ligated during RNA pol II ChIA-PET and sites linked by Cicero prompted us to investigate whether they might also be associated with other DNA binding protein complexes. We performed a search for known sequence motifs within each site that could accurately predict which other sites, if any, Cicero would link to it. Promoters with DNA binding motifs for core myogenic transcription factors were significantly more likely to be connected to an opening distal site than promoters without them (co-accessibility score > 0.25). For example, promoters containing at least one MYOD, MYOG or MYF6 motif were 4.0-fold more likely to be connected to an opening distal site than promoters with none of these motifs (p = 9.6 x 10^-260^; likelihood ratio test for logistic regression model), and similarly, promoters with at least one MEF2 family motif were 2.5-fold more likely to be connected to an opening distal site (p = 3.4 x 10^-81^).

We hypothesized that these correlations between promoter motifs and dynamic changes in their linked distal sites resulted from physical, transcription factor-mediated interactions. To explore this further, we focused on promoters linked to exactly one dynamically accessible distal site (co-accessibility score > 0.05). We enumerated the motifs present in each such promoter-distal site pair, and then used the Graphical LASSO to identify pairs of motifs wherein the presence of a motif in the promoter was predictive of the presence of the paired motif in the dynamically accessible distal site (see Methods).

This analysis uncovered a number of motif pairs corresponding to transcription factors known to physically interact. For example, opening distal elements were significantly more likely to have a MEF2, MEIS1 or RUNX1 motif if they were linked to a promoter that included a MYOD motif (**Supplemental Figure 4**). These enrichments were generally reciprocal, *i.e.* holding true regardless of which motif was in a promoter and which was distal. Myogenic regulatory factors (MRFs) interact physically with the MEF2, MEIS1, and RUNX1 proteins(Knoepfler et al., 1999; Molkentin et al., 1995; Philipot et al., 2010). Indeed, in their report that physical interactions between MEF2 and MYOD greatly enhance target gene expression in co-transfected fibroblasts, Olson and colleagues first proposed that such interactions might establish a loop between distally located binding sites(Molkentin et al., 1995). The correlations between dynamic distal and promoter sites detected in single cell chromatin accessibility data are consistent with physical interactions between them, mediated by bound transcription factors.

### MYOD coordinates histone modifications in cohorts of co-accessible sites

The physical proximity of co-accessible sites suggested that recruitment of histone-modifying enzymes to one site might nucleate changes in others. Indeed, pairs of sites were more likely to be undergoing significant, concordant gains in H3K27ac if they were linked by Cicero (**Figure 5A**). Sites that themselves exhibited static accessibility, but were linked to a dynamic opening site, showed strong gains in H3K27ac, while static sites that were linked to dynamic, closing sites showed strong losses (**Figure 5B**). The gains in acetylation might be driven by *de novo* binding of MYOD at the opening site followed by recruitment of a histone acetyltransferase (*e.g.* p300). Supporting this, of the 1,769 sites with significant gains in H3K27ac but no change in accessibility, only 823 (47%) were bound by MYOD in myotubes. However, 97% were linked with positive co-accessibility to a MYOD-bound site. Similarly, static, deacetylated sites either lost MYOD binding themselves or were linked to another site that did so (**Figure 5C**).

**Figure 5.**
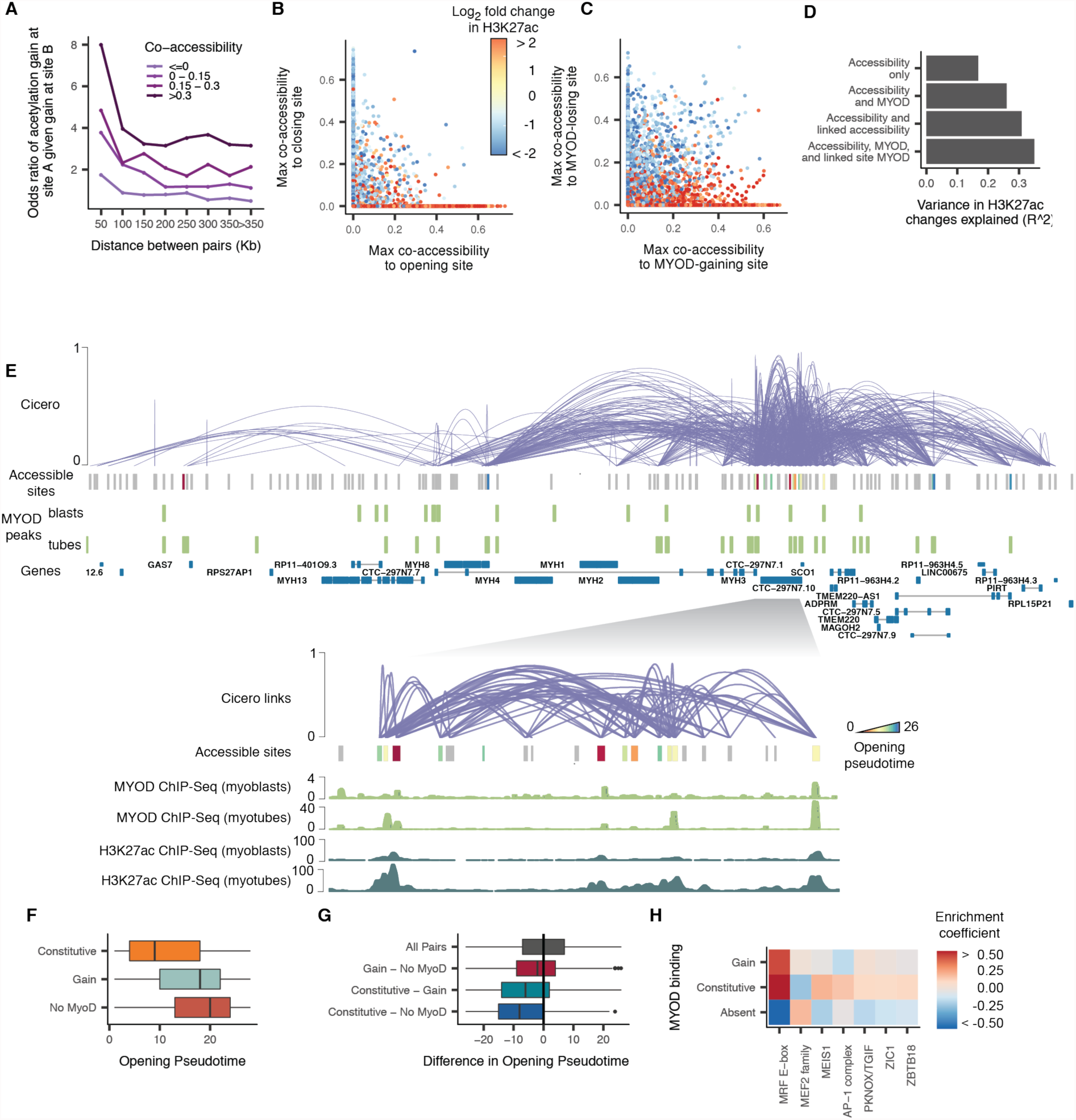
Co-accessible DNA elements linked by Cicero are epigenetically co-modified. **A**) Odds ratio of a site gaining H3K27ac during myoblast differentiation, given that it is linked to a site that is doing so. Color indicates the strength of the Cicero co-accessibility links. The lightest color indicates pairs of sites that are unlinked by Cicero. **B**) Correspondence between a statically accessible site’s gain or loss of H3K27ac and its maximum co-accessibility score to a site that is opening (x axis) or closing (y axis). Sites that are not linked to an opening or closing site are drawn at x = 0 or y = 0, respectively. **C**) Similar to panel B, but describing the correspondence between a site’s gain or loss of H3K27ac and its maximum co-accessibility score to a site that is gaining or losing MYOD. **D**) The variance explained in a series of linear regression models in which the (Gaussian) response is the log2 fold change in H3K27ac level of each DNA element and the predictors are whether that site is opening, closing, or static, whether it undergoes gains or loses in MYOD binding, and whether it is linked to neighbors that are doing so. See Methods for full details on model specifications. **E**) The Cicero map for the 755kb region surrounding *MYH3* along with called MYOD ChIP-seq peaks from (Cao et al., 2010). Sites opening in accessibility are colored by their opening pseudotime (see Methods), sites that are not opening in accessibility are shown in grey. The inset shows the 60kb region surrounding *MYH3* along with MYOD ChIP-seq and H3K27ac ChIP-seq signal tracks from (Cao et al., 2010) and (The ENCODE Project Consortium, 2012). **F**) Opening pseudotimes for all opening sites, subdivided by whether MYOD is bound in myoblasts and myotubes, myotubes alone, or neither. **G**) The difference in opening pseudotimes between pairs of linked DNA elements. The pairs are grouped based on whether one or both sites is constitutively bound by MYOD. For example, the bottom boxplot shows that sites that are constitutively bound by MYOD open earlier than the unbound sites to which they are linked. **H**) Transcription factor binding motifs selected by an elastic net regression (alpha = 0.5), with a response encoding the MYOD binding status of each site.

To quantify the predictive power of MYOD binding dynamics, we constructed a linear regression model that aimed to explain changes in histone modifications at each accessible site based on whether MYOD binding was gained, lost, maintained, or absent throughout differentiation. Sites gaining MYOD were significantly more likely to gain a variety of histone marks associated with activated regulatory elements, including H3K27ac and H2A.Z (**Figure 5D; Supplemental Figure 5**). However, this model explained only 26% of variation in H3K27 acetylation. When we expanded it to also include each site’s maximum co-accessibility score to a linked site with a gain in MYOD binding, the model improved to explain 35% of the variation, a 1.35-fold increase. Similar analyses of other marks showed comparable gains in explanatory power when Cicero linkages were included, supporting the view that sites predicted to form chromatin hubs by Cicero are coordinately targeted by histone modifying enzymes.

Although recruitment of MYOD to chromatin hubs explains a substantial proportion of their H3K27ac dynamics, it remained unclear whether gains in MYOD were concentrated in a few CCANs (*i.e.* hubs) as opposed to being widely distributed. Of the 2,561 hubs that contained gene promoters, 68% contained at least one site undergoing a gain or loss in MYOD binding, with some containing more than one MYOD binding site. For the subset of 462 hubs with a differentially expressed gene, 88% contained at least one site changing in MYOD binding. For example, within the single hub enclosing myosin heavy chain isoforms 1, 2, 3, 4, 8, and 13 along with numerous other genes, 18 sites underwent significant increases in accessibility. Of these, 13 were bound by MYOD in myotubes (**Figure 5E**). Interestingly, however, two sites very near *MYH3* (marked with asterixes) opened substantially earlier in pseudotime than the others and were bound by MYOD in myoblasts as well.

We wondered more generally whether sites bound by MYOD in myoblasts and throughout differentiation opened earlier than sites that gained MYOD binding during differentiation. A changepoint analysis using the PELT algorithm (Killick et al., 2012) revealed that DNA elements bound by MYOD throughout differentiation opened significantly earlier than those that gained MYOD (Mann-Whitney test p-value 4.2e-89) or were never bound by it (Mann-Whitney test p-value 1.0e-133) (**Figure 5F**). Moreover, rather than being enriched in whole hubs that open early as a group, constitutively MYOD-bound sites opened significantly earlier than sites linked to them that either gained MYOD (two-sided paired Student’s t-test p-value = 6.0e-120) or were never bound by it (two-sided paired Student’s t-test p-value = 6.4e-138) (**Figure 5G**). These sites with constitutively bound MYOD were enriched for some TF binding motifs that were not enriched in sites that gained MYOD during differentiation. Specifically, while both classes of MYOD-bound sites contained MRF E-boxes, only the constitutively MYOD-bound sites were enriched for the MEIS1 motif (**Figure 5H**). While 14% of sites within chromatin hubs contained MEIS1 motifs, 30% of constitutively MYOD-bound, dynamically opening sites in chromatin hubs contained them. Altogether, sites with MEIS1 motifs were linked to 66% of dynamically opening sites, and 54% of the sites that gained H3K27ac, compared with only 14% of sites genomewide (co-accessibility > 0.25). Murine Meis1, in conjunction with Pbx1, has been reported to act as a complex required for the MYOD-mediated activation of the myogenin promoter, and mutations in MYOD that prevent interaction with PBX resulted in loss of binding and regulation of roughly 10% of its sites and regulated genes(Berkes et al., 2004; Fong et al., 2015). Our results suggest that MEIS1 recruitment of MYOD may be pervasive throughout the genome, and could nucleate activation of other sites within a chromatin hub.

### Sequence features of active chromatin hubs predict gene regulation

We found that differentially expressed genes showed greater correlation in expression as a function of their Cicero co-accessibility score (**Supplemental Figure 6**). We next asked whether Cicero co-accessibility could be used to predict changes in gene expression. We first devised a linear regression model that aimed to predict changes in either expression or a variety of epigenetic marks associated with gene activation (**Figure 6A**). However, chromatin accessibility at distal sites might be correlated with epigenetic marks or expression with many genes in the surrounding region, potentially yielding a false impression of the predictive power of a model trained using it. To avoid this, we trained our model to predict gene activity using sequence features alone.

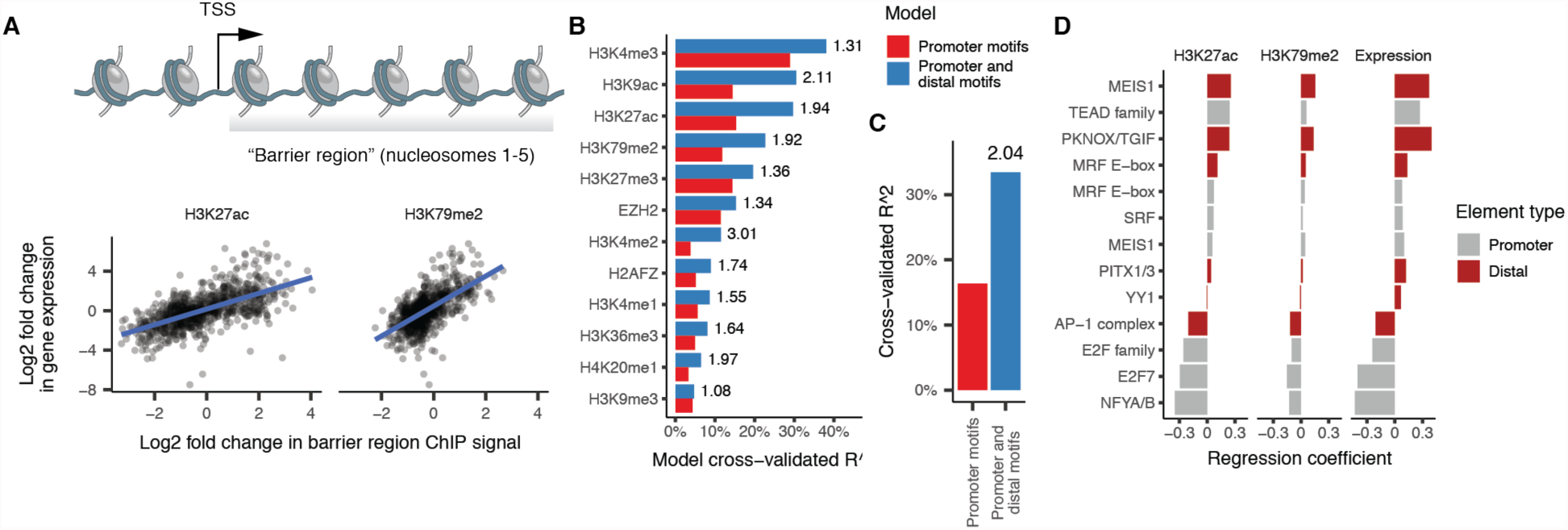
**Figure 6**. Chromatin dynamics at distal DNA elements predicts gene regulation. **A**) Changes in histone acetylation in the first 1 kb downstream of each gene’s TSS, corresponding to the “barrier” to RNA pol II elongation posed by nucleosomes, are correlated with changes in its expression. **B**) Two regression models predict changes in the histone marks deposited throughout each gene’s barrier region. The first model predicts changes on the basis of transcription factor binding motifs in gene promoters. The second model adds variables encoding the strength of co-accessibility with linked sites containing the motif. See Methods for details on the various models. Adjusted R^2 is computed as the fraction of null deviance explained. The number to the right of each bar indicates the ratio of variance explained between the first and second model. **C**) Similar to panel B, with changes in expression as the response. **D**) Coefficients from the model incorporating sequence at distal sites for each motif surviving model selection via elastic net. Note that the model considers each motif twice: once at promoters and again at distal sites, and both can be selected by elastic net.

Our first model takes as input a binary map of the transcription factor binding motifs present at the promoter upstream of each TSS. We then train it to predict how much of a gene’s observed expression change is attributable to each TF motif using elastic net regression and 50-fold cross-validation (Methods). The promoter-based model explained only 16% of the variance in expression and performed similarly in predicting a panel of histone marks (**Figure 6B-C**).

We then augmented the model with TF motifs at distal sites linked to the promoter(s) of a TSS by Cicero. We computed, for each promoter and for each sequence motif, the sum of the co-accessibility between it and any distal element that contained the given sequence. That is, the model associated each TSS with a co-accessibility score for each TF motif, taking the sum of the scores if a TSS was linked to more than one distal site carrying it. This augmented model markedly improved our ability to predict changes in expression, attributing 33% of the variance to motifs alone, a 2.04-fold increase (**Figure 6C**). This model similarly explained more variance of both activating marks such as H3K27ac and H3K9ac as well as those associated with silencing such as H3K27me3 (**Figure 6B**). The TF motifs identified by the model included the MRF E-box, the MADS box bound by MEF2 family proteins, the MEIS1 binding site, which were associated with gene upregulation, along with motifs for factors that drive cell proliferation such as AP-1, which were linked to downregulation. Importantly, motifs for the MRF family and MEIS1 survived the elastic net feature selection procedure at both promoters and distal sites, indicating that distal sites provide complementary, rather than redundant, information to the model (**Figure 6D**). Thus, when tasked with predicting which factors are important for gene regulation, our regression identified the major myogenic transcription factors using only the sequences in sites linked together by Cicero.

## Discussion

Despite their paramount importance for interpreting GWAS as well as for our basic understanding of gene regulation, we still lack comprehensive maps that link distal regulatory sequences to their target genes. There are several reasons for this. First, current methods for constructing genome-wide regulatory maps require input data collected from diverse tissues and cell lines. Second, analyzing these data poses a major statistical challenge, especially in the presence of batch effects and other technical features of measurements collected at different times by different labs. Third, a map produced by integrating data from many tissues and cell types might capture interactions common to many cell types at the expense of cell-type specific interactions.

Here, we describe Cicero, which constructs cis-regulatory maps from single cell chromatin accessibility data such as is generated by sci-ATAC-seq. Cicero exploits the fact that patterns of co-accessibility between regulatory elements located in cis derive in part from physical interactions between the transcription factors that mediate gene regulation. Maps obtained through the application of Cicero may advance our quantitative understanding of the logic of gene regulation in eukaryotic genomes, while also advancing our ability to identify the target genes of noncoding GWAS signals.

Pseudotemporal ordering of chromatin accessibility profiles from differentiating myoblasts, a classic model system of vertebrate developmental gene regulation, revealed dynamic changes in thousands of DNA elements. Although changes in promoter accessibility proved to be a poor predictor of gene expression, distal sites linked to genes by Cicero markedly improved power to predict gene regulation. In this study, we used easy-to-interpret linear regression techniques to investigate the role of distal elements in regulating target genes, but more sophisticated machine learning approaches might make better use of Cicero’s regulatory maps.

Taken together, our analyses show that the cis regulatory elements linked by Cicero meet the definition of chromatin hubs: they are physically close in the nucleus, their histone marks change in a coordinated fashion, and their interactions are likely mediated by a common set of DNA binding proteins, including lineage-specific transcription factors. For myogenesis, our results support a model of gene activation in which a subset of “precocious” enhancers recruit chromatin remodeling enzymes and other epigenetic modifiers to the hub, which then mediate increases in accessibility of other binding sites (**Figure 7**). In differentiating myoblasts, MYOD is widely understood to recruit the BAF complex and p300/PCAF to activate enhancers of muscle genes (Serra et al., 2007; Simone et al., 2004). Although the role of MYOD in recruiting these is well appreciated, how MYOD is itself recruited to its binding sites in their inaccessible state is less clear. Our analyses of accessibility dynamics of regulatory elements within individual chromatin hubs showed that rather than simultaneously opening as a group, a subset of sites often open much earlier than the rest of the hub. These early-opening sites are distinguished by enrichment for MEIS1 motifs and the presence of MYOD throughout myoblast differentiation. Meis1 has previously been reported to tether Myod to the inactive myogenin promoter prior to the onset of differentiation, and is required for myogenin activation and chromatin remodeling that permits the binding of MYOD to nearby MRF E-boxes that were previously inaccessible (Berkes et al., 2004; Maves et al., 2007; de la Serna et al., 2005). Whether Meis1/Pbx1 acts to tether MYOD to inactive chromatin more generally throughout the genome has remained an open question, however, MYOD mutations that prevent interaction with PBX prevented binding at approximately 10% of MYOD bound regions (Fong et al., 2015), suggesting the possibility of a broad role in the myogenic program. Our analyses, which show the enrichment of MEIS1 motifs in sites bound by MYOD prior to the onset of differentiation, but not sites bound later, suggest that MEIS1 (and likely its co-factor PBX1) serve as the initial recruitment sites for epigenetic remodeling enzymes. Binding of p300 to MEIS1/PBX1-tethered MYOD could then acetylate histones at all DNA elements physically nearby in the chromatin hub. This model may help explain the pervasive gains and losses of histone acetylation throughout the accessible genome, despite the comparatively much smaller number of differentially accessible or MYOD-bound elements. This model is also consistent with a recent study by Hilton *et al.* which showed that directly recruiting a Cas9/p300 fusion protein to distal enhancers upstream of the MYOD, OCT4, and globin loci increased expression and induced histone acetylation at their respective promoters in 293T cells (Hilton et al., 2015).

**Figure 7.**
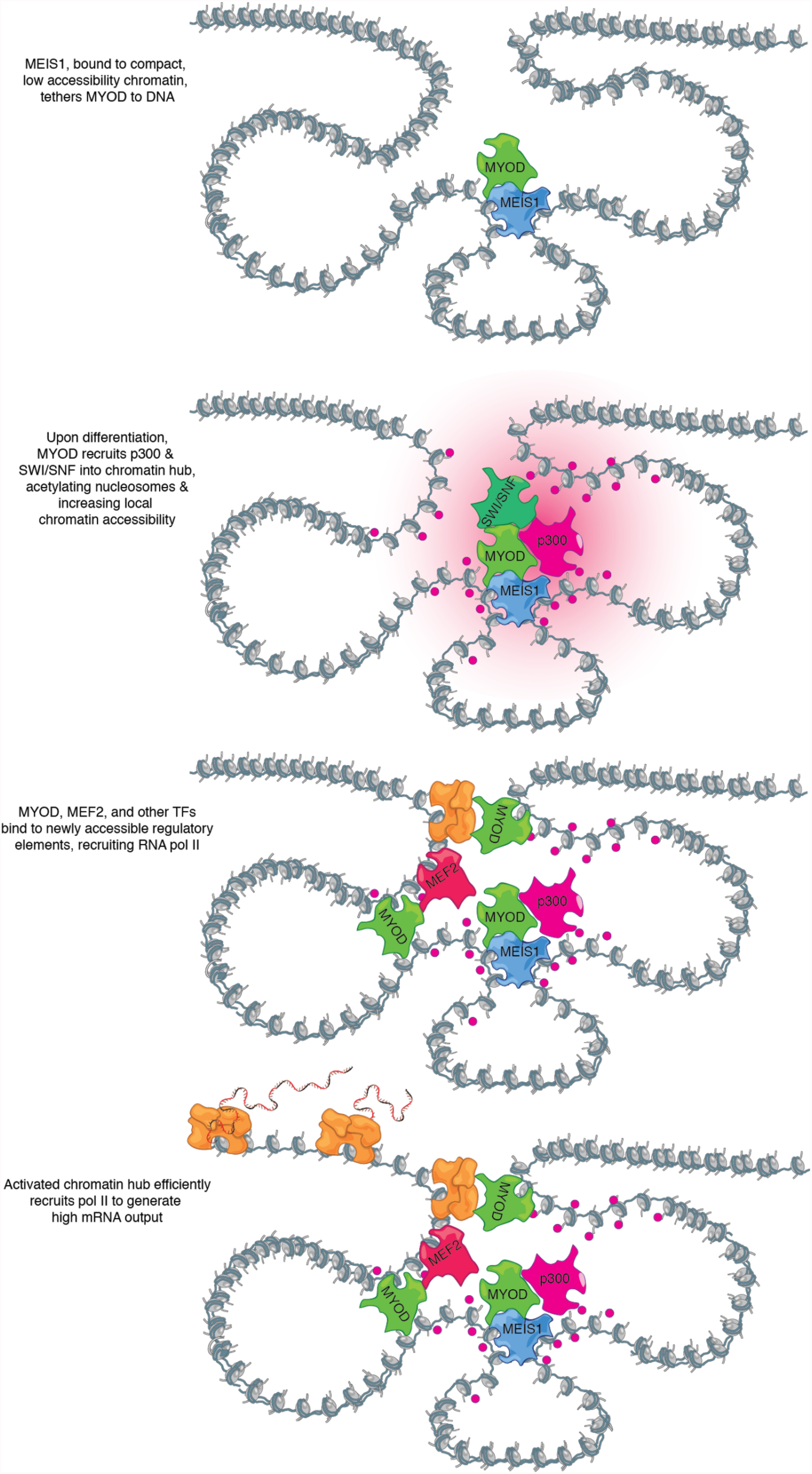
A model of how chromatin hub activation could be nucleated by a subset of “precociously” opening DNA elements within it. Such sites are occupied by transcription factors competent to bind relatively closed, inactive DNA elements, such as MEIS1, which may tether less competent factors such as MYOD to the hub. Subsequent recruitment of p300 and the BAF complex, possibly through intermediary factors (e.g. MYOD), leads to remodeling and acetylation of histones throughout other DNA elements nearby in the hub. These newly available sites are then bound by other transcriptional activators (e.g. MEF2), leading to the recruitment of Pol II. Moreover, acetylation of the histones downstream of assembled pre-initiation complexes reduces the barrier they pose to elongation, enhancing efficient transcription of genes within the hub.

Cicero provides an effective means of linking regulatory elements to their target genes in a tissue or cell type of interest using data from a single experiment. The chromatin hubs that it defines will facilitate the construction of quantitative models of epigenetic and gene expression dynamics, as well as the identification of genes whose dysregulation underlies GWAS associations. As the field pursues organism-scale cell atlases that comprehensively define each cell type and its molecular profile, such regulatory maps will be essential for understanding the epigenetic basis of each cell type’s gene expression program, both in health and disease.

## Methods

### Cell culture (HSMM and GM12878)

HSMM derived from quadriceps biopsy (Lonza, catalog #CC-2580, lot #257130: healthy, age 17, female, of European ancestry, body mass index 19; cells were used within 5 passages of purchase) were cultured in skeletal muscle growth media (GM) using the SKGM-2 BulletKit (Lonza). The cells and differentiation protocol are those from Trapnell *et al.* (2014). Cells were seeded in 15 cm dishes, media being replenished every 48 hours and cells were seeded in 15 cm dishes, cells allowed to reach 80-90% confluence. Differentiation was induced at time 0 via a switch to differentiation medium (DM) composed of alpha-mem (Thermo Fisher Scientific) and 2% horse serum. Cells in GM (time 0) or DM were then harvested at the specified times and processed as described below. HSMM tested negative for mycoplasma contamination within 6 months of the experiment.

GM12878 (purchased from Coriell Cell Repository) was cultured in RPMI 1640 medium (Gibco 11875) supplemented with 15% FBS, 100U/ml penicillin and 100 μg/ml streptomycin. Cells were cultured in an incubator at 37C with 5% CO2 and were split to a density of 300,000 cells/ml three times a week.

### Sci-ATAC-seq library construction

We prepared sci-ATAC-seq libraries using an improved version of the original protocol (Cusanovich et al, 2017, *submitted*).

### Defining accessible sites

To define peaks of accessibility across all sites, we used the MACS (version 2.1.0) (Zhang et al., 2008) peak caller. Cells with fewer than 1,000 reads were filtered, and reads from repeat-masked regions of the genome were excluded from peak-calling. Promoter peaks were further defined as the union of the annotated transcription start site (TSS) (Gencode V17) minus 500 base pairs, and MACS defined peaks upstream of the TSS. Cells were determined to be accessible at a given peak if a read from that cell overlapped the peak.

560 barcodes from the HSMM dataset and 100 barcodes from the GM12878/HL60 dataset with a high percentage of peaks with more than 2 reads mapping to them were excluded as potential doublets.

For the GM12878 and HL60 mixed dataset, preliminary peaks were called by MACS and used to separate the cell types using multi-dimensional scaling by Jaccard distance. The subset of reads from GM12878 cells was then used to recall peaks for GM12878 as above.

### Pseudotemporal ordering

For the HSMM dataset, contaminating interstitial fibroblasts were removed in silico based on the absence of promoter accessibility in any of several known muscle markers (MYOG, MYOD1, DMD, TNNT1, MYH1, TPM2). In addition, cells with fewer than 1,000 accessible sites were excluded due to low assay efficiency. Finally, peaks present in less than 1% of cells were excluded during pseudotemporal ordering steps.

Despite improvements to the sci-ATAC-seq protocol that delivered a substantial increase in the number of sites detected per cell, sci-ATAC-seq data remains zero-inflated. The quality and efficiency of transposition, which varies between cells and across batches, is likely to be a major technical source of variation in the data. Simple dimensionality reduction techniques such as MDS show that a poorly-assayed cell is often more similar to other poorly-assayed cells of a different type than to well-assayed cells of the same type. In order to accurately group cells with similar chromatin accessibility profiles, we first clustered peaks that were within 1 kb and summed the reads overlapping them to create an integer-valued count matrix *M*.

To order the cells by progress through differentiation, we determined which aggregated peaks were relevant to the HSMM time course by fitting the following model:

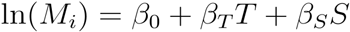

Where *M_i_* is the mean of a negative binomially-distributed random variable for the number of reads overlapping the aggregate region *i*, *T* encodes the times at which each cell was harvested and *S* is the total number of accessible sites in each cell. We compared this full model to the reduced model:

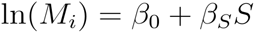

by likelihood ratio test. Sites determined by this method to be time dependent and which were accessible in less than 10% of cells were then used to reconstruct the pseudotime trajectory using Monocle 2 (parameters ncenter and param.gamma set to 100, see (Qiu et al., 2017)). To remove any bias created by different assay efficiency in different cells, total sites accessible was included as a covariate in the tree reconstruction. Each cell was assigned a pseudotime value based on its position along the trajectory tree. Cells that mapped to the F_2_ branch were excluded from downstream analysis.

### Differential accessibility analysis

When testing for differential accessibility across cells at a particular site, it is important to exclude technical variation due to differences in assay efficiency as discussed above. We first grouped cells at similar positions in pseudotime. We did this by k-means clustering along the pseudotime axis (k=10). These clusters were further subdivided such into groups containing at least 50 and no more than 100 cells. Next, we aggregated the binary accessibility profiles of the cells in each group into a matrix *A*, so that *A_ij_* contains the number of cells in group *j* for which DNA element *i* is accessible. The average pseudotime *ψ_j_* and average overall cell-wise accessibility *S_j_* for cells in each group *j* were preserved for use during differential analysis.

To determine which peaks of accessibility were changing across pseudotime, we fit the following model to the binned data:

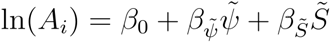

Where *A_i_* is the mean of a negative-binomial valued random variable of cells in which site is *i* accessible, and the tilde above *ψ* and *S* indicates that these predictors are smoothed with natural splines during fitting. This model was compared to the reduced model:

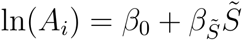

by the likelihood ratio test. Peaks with an adjusted p-value of less than 0.05 were determined to be dynamic across pseudotime.

#### Gene set enrichment analysis

Gene set enrichment analysis was conducted using the R package piano (Väremo et al., 2013) using a hypergeometric test. We tested against the Human GO Biological Processes gene set from (Merico et al., 2010).

### Cicero

Cicero aims to identify all pairs of co-accessible sites. The algorithm takes as input a matrix of *m* by *n* binary accessibility values *A*, where *A_mn_* is zero if no read was observed to overlap peak in cell and one otherwise. The algorithm also requires either a pseudotemporal ordering of the cells along a developmental trajectory (e.g. with Monocle 2) or the coordinates of the cells in some sufficiently low dimensional space (e.g. a t-SNE map) that the cells can be readily clustered. The algorithm then executes the following steps, which are detailed in the sections below: first, groups of highly similar cells are sampled using the clustering or pseudotemporal ordering, and their binary profiles are aggregated into integer counts. Second, these counts are optionally adjusted for user-defined technical factors, such as experimental batch. Third, Cicero computes the raw covariances between each pair of sites within overlapping windows of the genome. Within each window, Cicero estimates a regularized correlation matrix using the graphical LASSO, penalizing pairs of distant sites more than proximal sites. Fourth, these overlapping covariance matrices are “reconciled” to produce a single estimate of the correlation in accessibility across groups of cells. These correlation scores are reported to the user, who can extract modules of sites that are connected in co-accessibility networks by first specifying a minimum correlation score and then using the Louvain community detection algorithm on the subgraph induced by excluding edges below this score.

#### Grouping cells

In principle, Cicero could analyze the sample covariance computed between the vectors *x_i_* and *x_j_* of binary values encoding accessibility across cells for a pair of *i* sites and *j*. However, rather than working with the binary data directly, Cicero groups similar cells and aggregates their binary accessibility profiles into integer count vectors that are easier to work with in downstream steps. Under the grouping discussed below, the number of cells in which a particular site is accessible can be modeled with a binomial distribution or, for sufficiently large groups, the corresponding Gaussian approximation. Modeling grouped accessibility counts as normally distributed allows Cicero to easily adjust them for arbitrary technical covariates by simply fitting a linear model and taking the residuals with respect to it as the adjusted accessibility score for each group of cells.

In order to control for technical variation as discussed above, Cicero operates on a grouped cell count matrix, *C*. *C* is constructed by first mapping cells into 2 dimensions by either Monocle 2 or tSNE. Within this space, Cicero constructs a k-nearest neighbor graph, via the the FNN package, which is based on KD-trees and is highly efficient and scales to large numbers of cells. Cicero then selects *d* random cells, and their *k* nearest neighbors are grouped. Accessibility counts are then summed across all cells in a group to create count matrix *C*.

#### Adjusting accessibility counts for technical factors

To normalize for variations in assay efficiency across groups, matrix *C* is divided by a group wise scaling factor (computed using the standard Monocle 2 method for library size calculations (`estimateSizeFactors ()`) to create an adjusted accessibility matrix *R*. Because the entries of *C* are integer counts that can reasonably be approximated by Gaussian distributions, this matrix can be readily adjusted for arbitrary technical covariates (e. g. using the Limma package’s `removeBatchEffect ()` function). In this study we did not adjust for factors beyond library size.

#### Computing co-accessibility scores between sites

Cicero next analyzes the covariance structure of the adjusted accessibilities in *R*. Given enough data, Cicero could in principle simply compute the raw covariance matrix *U*. However, because the number of possible pairs of sites is far larger than the number of groups of cells, Cicero uses the Graphical Lasso to compute a regularized covariance matrix to capture the co-accessibility structure of the sites. The Graphical LASSO computes the inverse of the sample covariance matrix, which encodes the partial correlations between those variables as well as the regularized covariance matrix (Friedman et al., 2008). These constitute a statistically parsimonious description of the correlation structure in the data: informally, two variables are partially correlated when they remain correlated even after the effects of all other variables in the matrix are excluded. The Graphical LASSO expects a small fraction of the possible pairs of variables to be partially correlated, preferring to select a sparse inverse covariance matrix over a dense one that fits the data equally well. Those pairs of sites that lack sufficient partial correlation to be worth the penalty term are assigned zero partial correlation in the inverse covariance matrix reported by Graphical LASSO. Formally, Cicero uses Graphical LASSO to maximize:

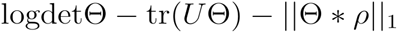

Where Θ is the inverse covariance matrix capturing the conditional dependence structure of *p* accessible sites, and *U* is the sample covariance matrix computed from their values in. In order to ensure stability of GLASSO, which can hang on poorly conditioned input, we add a small conditioning constant of 1e-4 to the diagonal of *U* prior to running it. The matrix *ρ* contains penalties that are used to independently penalize the covariances between pairs of sites, and denotes component-wise multiplication.

In Cicero, we aim to find local cis-regulatory interactions, rather than global covariance structure that might be expected due to overall cell state. To achieve this, we set each penalty term in *ρ* such that peaks closer in genomic distance had a lower penalty term. Specifically, we used the following equation to determine *ρ*:

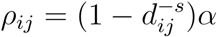

Where *d_ij_* is the distance in the genome (in kilobases) between sites *i* and *j* and *s* is a constant that captures the power-law distribution of contact frequencies between different locations in the genome as a function of their linear distance. A complete discussion of the various polymer models of DNA packed into the nucleus is beyond the scope of this paper, but we refer readers to (Dekker et al., 2013) for a discussion of justifiable values for *s*. We use a value of 0.75 by default in Cicero, which corresponds to the “tension globule” polymer model of DNA (Sanborn et al., 2015). The scaling parameter *α* controls the distance at which Cicero expects no meaningful cis-regulatory contacts, and its value is calculated automatically from the data. To calculate *α*, Cicero selects 200 random 500 kb genomic windows, and determines the minimum *α* value such that no more than 10% of pairs of sites at a distance greater than 250 kb (a user-adjustable value) had non-zero entries in Θ. The mean of these values of *α* is then used to set the penalties for the whole genome. Cicero then applies Graphical LASSO to overlapping 500 kb windows of the genome (windows are spaced by 250 kb such that each region is covered by two windows).

#### Reconciling overlapping local co-accessibility maps

Cicero calculates correlation values (co-accessibility scores) from the resulting estimated sparse covariance matrix for each pair of peaks within 500 kb of each other. Because the genomic windows are overlapping, the majority of pairs of peaks have two calculations of co-accessibility. To consolidate these sites and create a genome-wide map of the accessible regulome, Cicero considers the co-accessibility scores for each pair of peaks to determine if they are in qualitative agreement (both calculated scores in the same direction). The qualitative agreement in our two test datasets were both >95%. Pairs of peaks not in qualitative agreement are considered undetermined. For peaks in qualitative agreement, the mean score of the two values is assigned.

#### Extracting cis-co-accessibility networks (CCANs)

Positive Cicero co-accessibility scores indicate that a pair of peaks is connected, with the magnitude of the co-accessibility corresponding to Cicero’s confidence in the link. To identify hubs of co-accessibility, Cicero can create a graph where each node is a peak of accessibility, and edges are the co-accessibility scores above a user-defined threshold. Communities within this genome-wide graph can be found using the Louvain community finding algorithm. Cicero can then assign peaks to cis-coaccessibility networks (CCANs) based on these communities.

### Motif enrichment analysis

Transcription factor motifs from the JASPAR 2016 database (Mathelier et al., 2016) were located in the sci-ATAC-seq peaks using FIMO (Grant et al., 2011). Motifs for TFs not expressed at ≥ 2 transcripts per million in bulk RNA-seq (HSMM myoblasts or myotubes) were excluded from downstream analysis. Many TF motifs are similar or identical to each other. To prevent this correlation from confounding regression analyses, we clustered motifs into motif families. For each pair of motifs A and B, we computed the conditional probability that given motif A is called at a genomic location with a FIMO p-value < 2e-5 (a stringent threshold), an overlapping instance of motif B will be called at p < 1e-4 (a permissive threshold). We constructed an undirected graph of motifs where there is an edge between motifs A and B if P(B at p < 1e-4 | A at p < 2e-5) ≥ 0.5 or P(A at p < 1e-4 | B at p < 2e-5) ≥ 0.5. Edges in this graph are assigned weights equal to the greater of these two conditional probabilities minus 0.5. We clustered the motifs on this graph using Louvain clustering (Blondel et al., 2008) and manually assigned names to each cluster. For downstream regression analyses, a genomic location is considered to have an instance of a motif family if any motif in the family is called at that location at p < 5e-5 (an intermediate threshold).

To generate the motif co-accessibility networks shown in **Supplemental Figure 4**, we computed two sets of binary variables for each protein coding gene that had at least one sci-ATAC-seq peak in its promoter(s). The first set of variables are indicators of whether or not at least one instance of a motif family is present in any promoter peak for the gene. The second set of variables are indicators of whether or not at least one motif instance is present in any distal peak (excluding promoters of other genes) that is within the same Cicero CCAN (correlation score > 0.05) as the gene’s promoter(s). We constructed a matrix where rows are genes and columns are these two sets of motif indicator variables. This matrix was provided as input to the Graphical LASSO subject to the constraint that partial correlations between two promoter motif variables or two distal motif variables are fixed to zero. The regularization parameter ρ for the Graphical LASSO was set as the smallest value that could achieve an estimated false discovery rate (FDR, the proportion of truly-zero partial correlations that are estimated as non-zero) of less than 5%. The FDR for a given value of ρ was estimated by running the Graphical LASSO with that value of ρ on versions of the motif indicator matrix with the distal variables row-shuffled (essentially assigning each gene to a random other gene’s set of distal motifs) and counting the proportion of motif pairs that are assigned a non-zero partial correlation (ideally, all should be zero in a shuffled matrix).

In **Supplemental Figure 4**, an edge is drawn between a pair of motif families A and B if both 1) the partial correlation of the indicator variable for A being at a distal site to the indicator variable of B being at a linked promoter site is > 0.02, and 2) the same is true if B is in the distal position and A is in the promoter.

### Analysis of ChIA-PET and Hi-C data

To compare our Cicero connections to Hi-C data, we used publicly accessible GM12878 data (Rao et al., 2014) (GSE63525) at 5 kb resolution. The intrachromosomal raw contact matrices were normalized using the provided normalization vector obtained using the matrix balancing procedure of Knight and Ruiz and described by Rao et al. The normalized contact matrices were further transformed by dividing by the genome-wide model of interaction probability as a function of 1-dimensional genomic distance also described by Rao et al. By dividing by the expected contact probability based on distance, we were able to consider Hi-C interactions beyond those expected by linear genomic distance. To compare our data to Hi-C, we first assigned each peak to its appropriate 5 kb bin, and then considered the mean observed/expected Hi-C contact probability at various distances between bins connected by Cicero at various co-accessibility score cutoffs. If two bins were connected by multiple co-accessibility scores, the bin connection was categorized based on the largest score. As a comparison, we also calculated the mean contact probability between bins containing accessible sites with co-accessibility scores less than or equal to zero. Lastly, **Figure 4A** includes the mean observed/expected Hi-C contact probability across all somatic chromosomes.

As a second comparison dataset, we used publicly accessible GM12878 polII ChIA-PET data (Tang et al., 2015) (GSE72816). To compare these data to Cicero’s connections, we first looked for overlap between our peaks, and ChIA-PET anchors. Because ChIA-PET anchors often overlap each other, we first merged overlapping anchors to create comparable ChIA-PET “peaks”. We considered accessible peaks within 1 kb of ChIA-PET peaks to be overlapping. To generate **Figure 4C-E**, we considered the subset of ChIA-PET and Cicero connections where the peaks were present in both datasets.

### Analysis of ChIP-Seq data (MYOD and histone)

To compare our accessible peaks to the known myogenesis master regulator MyoD, we used publicly accessible MyoD ChIP-seq in human myoblast and human myotube(MacQuarrie et al., 2013) (GSE50413). We considered our peaks to be bound by MyoD if they overlapped one of the annotated MacQuarrie et al. ChIP-seq peaks.

To compare our accessible peaks to histone modifications, we used publicly accessible ENCODE datasets in HSMM and HSMMtube (The ENCODE Project Consortium, 2012) (ENCFF000BKV, ENCFF000BKW, ENCFF000BMB, ENCFF000BMD, ENCFF000BOI, ENCFF000BOJ, ENCFF000BPL, ENCFF000BPM). We counted both HSMM and HSMMtube histone ChIP-seq reads in each accessible peak. To determine whether sites were changing in accessibility between HSMM and HSMMtube, we used DESeq2 differential analysis (Love et al., 2014) (FDR < 5%). To determine whether the barrier regions of genes were differentially histone modified, we similarly used DESeq2 to compare the read counts in the first 1000 base pairs of each GENCODE v17 transcript in HSMM and HSMMtube datasets.

To compare agreement between H3K27 acetylation marks of peaks connected by Cicero, we divided the odds of a site gaining acetylation if it’s connected site gained acetylation by the odds of a site gaining acetylation is it is connected to a site that is not gaining acetylation (**Figure 5A**).

### Modeling H3K27 Acetylation Changes

To model changes in acetylation among linked sites (Figure 5D), we compared four linear regression models:

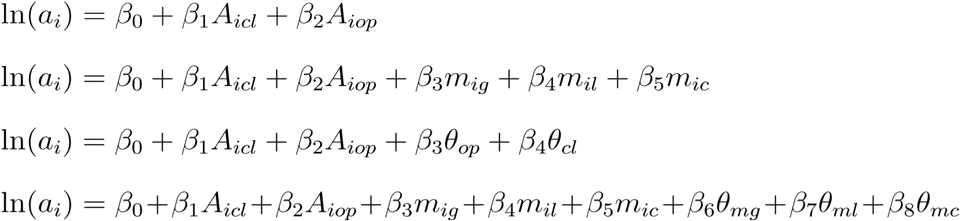

where *a_i_* is the log2 fold-change in H3K27 acetylation from myoblast to myotube at site *i*, *A_icl_* and *A_iop_* are indicator variables for whether site *i* is closing or opening across pseudotime, *m_ig_*, *m_il_* and *m_ic_* are indicator variables for whether site *i* is gaining, losing, or constitutively bound by MYOD from myoblast to myotube according to ChIP-seq, *θ_op_* and *θ_cl_* are the highest Cicero co-accessibility scores that connect site *i* to another opening or closing site respectively, and *θ_mg_*, *θ_ml_* and *θ_mc_* are the highest Cicero co-accessibility scores that connect site *i* to another MyoD gaining, MyoD losing or MyoD constitutive site. For each of the fitted models, we used elastic net regression (Zou and Hastie, 2005) to estimate the effect of each predictor.

Similarly, in Supplemental Figure 5, we predict the log2 fold-change in each of the 12 ENCODE histone mark ChIP-seq datasets described above using only indicator variables for whether a site is gaining losing or constitutively bound by MYOD, or using these variables and the highest Cicero co-accessibility scores connecting a site to an opening or closing site.

### Regression models for barrier region histone marks and gene expression

For each of the 12 ENCODE histone mark ChIP-seq datasets described previously, we fit two regression models that predict, for each transcription start site, the log fold change in the number of reads from the given ChIP-seq dataset that fall in the barrier region of that TSS (first 1000 bp downstream) for myotubes vs. myoblasts. We exclude TSSs that do not have a significantly different number of barrier region reads in myotubes vs. myoblasts for any of the 12 datasets (p > 0.01), leaving 6,205 TSS included in the model.

In the first set of models (“promoter motifs”), the features are a set of binary indicator variables that have value 1 if any promoter sci-ATAC-seq peak for the TSS has at least one instance of a motif from a given motif family. In the second set of models (“promoter and distal motifs”), the features are the promoter motif indicator variables plus a second set of real-valued variables that encode the presence of distal sequence motifs. For a given motif family and TSS, the corresponding distal motif variable has a value equal to the highest co-accessibility score from any promoter sci-ATAC-seq peak for that TSS to any connected distal peak that has at least one instance of a motif from the motif family. If no such distal peak exists (the motif is absent in all connected distal sites), the distal motif variable is assigned a value of 0. The models were trained using elastic net regression.

We additionally fit models with the same features (“promoter motifs” and “promoter and distal motifs”) to predict the expression of the subset of the above TSSs (n = 929), that were additionally expressed in at least 4 cells in scRNA-seq and which were predicted by smoothed average across pseudotime to be expressed at above 1 copy per cell at some pseudotime.

## Data Availability

Sci-ATAC-seq data will be made publicly available upon publication.

## Code Availability

We will release Cicero as an R package through Github and Bioconductor upon publication.

## Acknowledgements

We gratefully acknowledge Stephen Tapscott, William Noble, and Daniela Witten as well as members of the Shendure and Trapnell labs for their advice. This work was supported by the following funding: NIH grant U54 DK107979 to JS and CT; NIH grant DP2HD088158 to CT; NIH grants DP1HG007811 and R01HG006283 to JS; The Paul G. Allen Family Foundation to JS; JS is an Investigator of the Howard Hughes Medical Institute. CT is partly supported by an Alfred P. Sloan Foundation Research Fellowship. DAC was supported in part by T32HL007828 from the National Heart, Lung, and Blood Institute. JLM and AM are supported by NIH Genome Training Grant (5T32HG000035). HAP is supported by the National Science Foundation Graduate Research Fellowship under Grant No. (DGE-1256082).

---

**Supplemental Figure 1.**
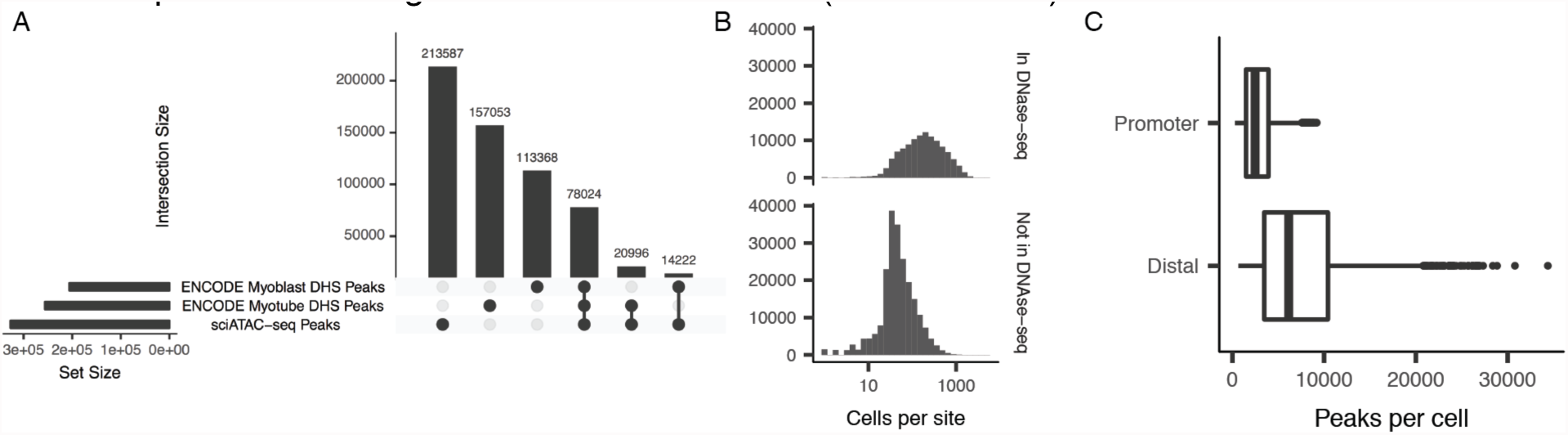
**A**) Overlap between ENCODE DnaseI hypersensitive sites (DHSs) and MACS-called sci-ATAC-seq peaks visualized as an UpSetR plot (Lex and Gehlenborg, 2014). Horizontal bars indicate the number of peaks in each data set. Vertical bars indicate the number of peaks in each compartment of the three-way Venn diagram. Black dots indicate which datasets are involved (i.e. intersected) in each vertical bar. **B**) Histogram of the number of cells with a read in each sci-ATAC-seq site, faceted by whether the site overlapped ENCODE DHS peaks in myoblasts or myotubes. **C)** Boxplot of the number of MACS-called sci-ATAC-seq peaks per cell. Promoter-proximal peaks are peaks intersecting the first 500 base pairs upstream of a transcription start site (see Methods). Distal peaks are all other peaks.

**Supplemental Figure 2.**
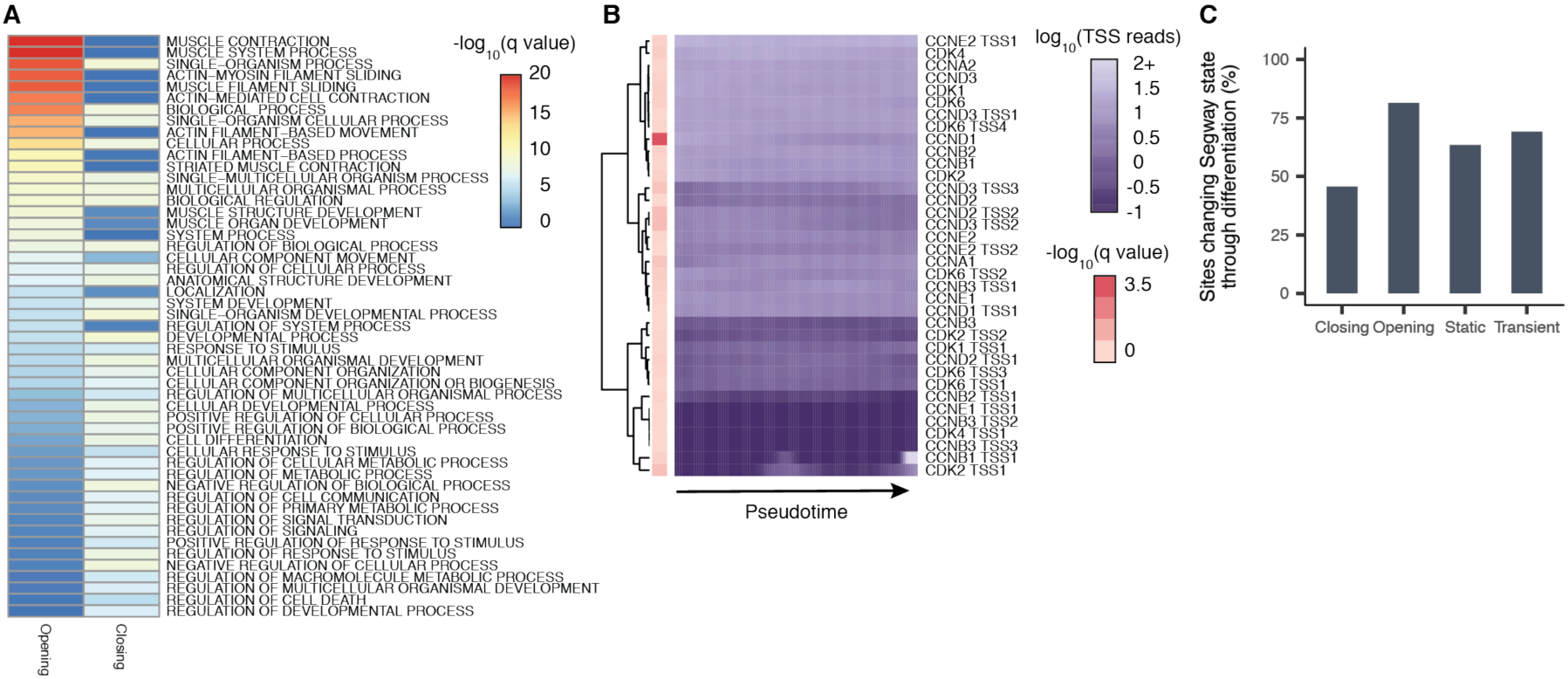
**A**) Gene set enrichment analysis of significantly opening and closing accessible sites. Adjusted p-values were computed using a hypergeometric test. Terms shown are all sites with an adjusted p-value < 1e-4 in either the opening set or the closing set. Color represent the -log10 adjusted p-value. Sites are ordered by the -log10 adjusted p-value of the opening set. **B**) Smoothed pseudotime-dependent accessibility curves, generated by a negative binomial regression of each for a set of selected cell cycle relevant genes. Each row indicates a different DNA element. Annotation column represents the -log10 adjusted p-value for the test of differential accessibility across pseudotime. For visualization, fitted curve range was capped at 100. **C**) Percent of dynamic and static sites with changing Segway state assignment from myoblast to myotube.

**Supplemental Figure 3.**
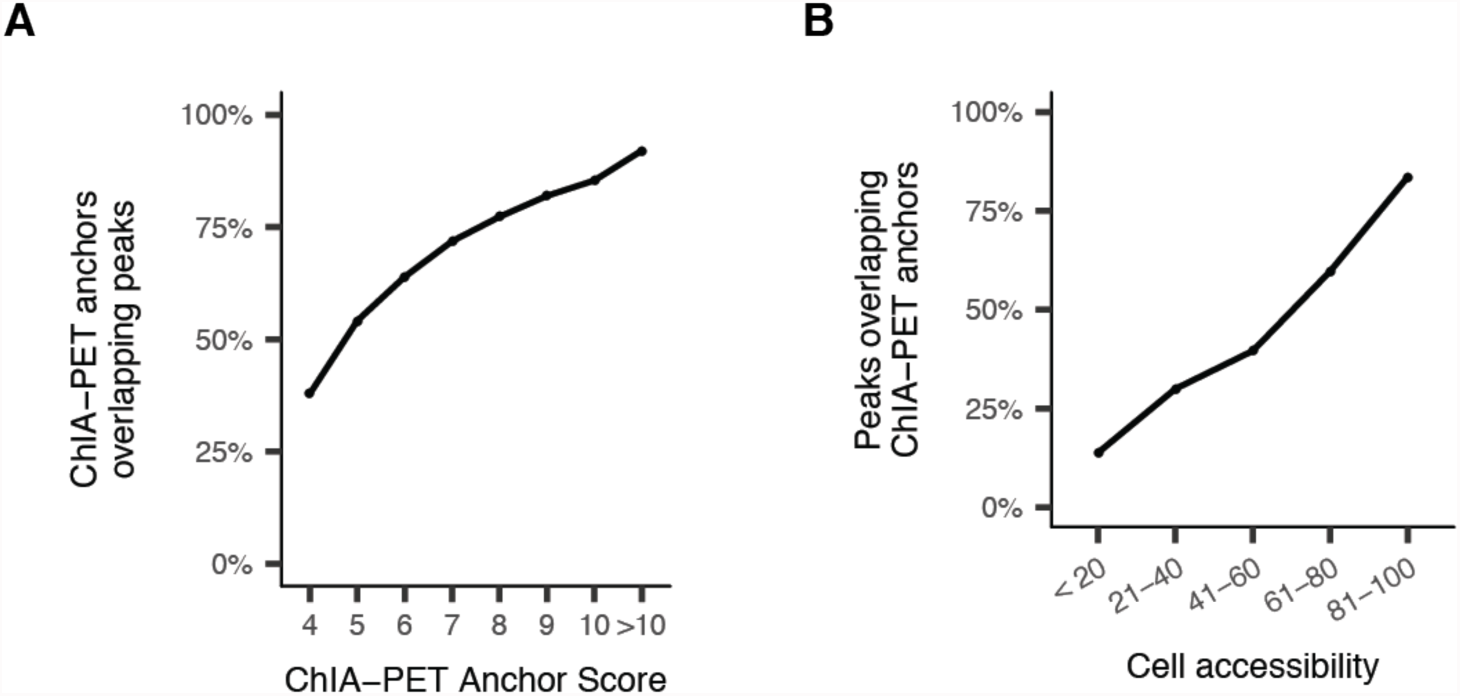
**A**) Percent of pol II ChIA-PET anchors within 1 kb of an sci-ATAC-seq peak as a function of ChIA-PET anchor score provided by Tang et. al. (2015). **B**) Percent of sci-ATAC-seq peaks within 1 kb of pol II ChIA - P ET anchors as a function of overall cell accessibility (number of cells where the peak is accessible).

**Supplemental Figure 4.**
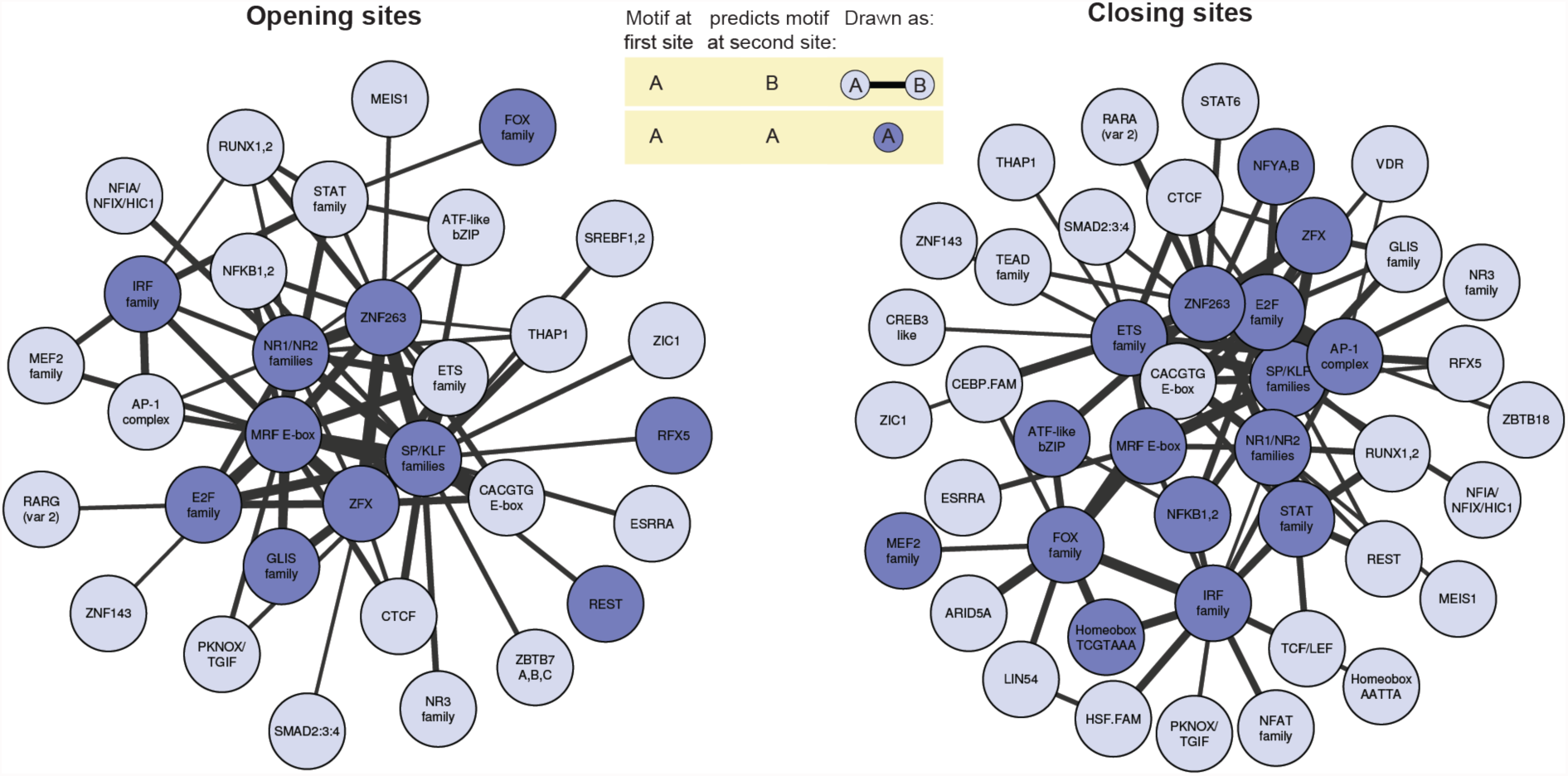
Motifs in accessible sites predict motif content of Cicero-linked sites. The network summarizes a graphical model that captures how occurrences of motifs in pairs of sites predict whether they are connected. Each motif is connected to the motifs it suggests will exist in one or more connected sites. A motif that predicts itself in a connected site is shown in dark blue. If motif “A” at a distal site predicts that “B” will be found at a promoter, and symmetrically “B” at a distal site suggests “A” will be found at a promoter, they are connected with a black line, with a width proportional to the strength of the co-accessibility. Asymmetric motif relationships are not shown.

**Supplemental Figure 5.**
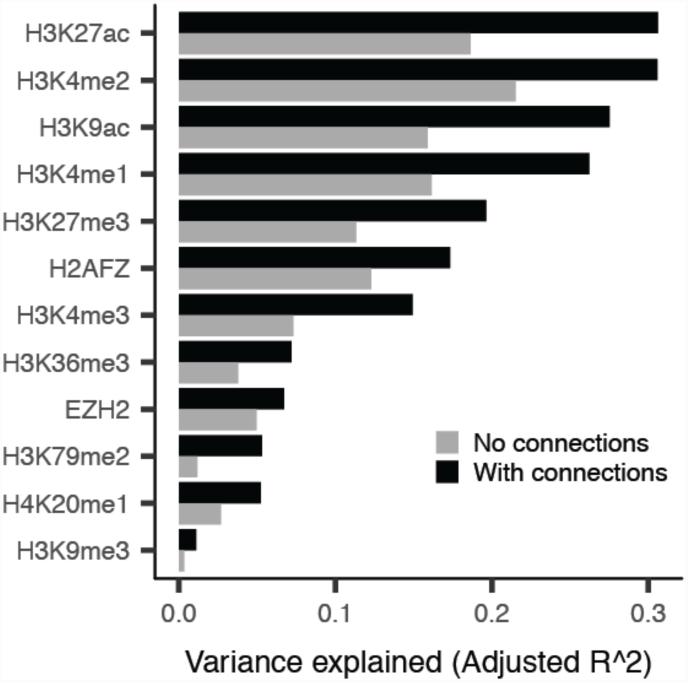
Variance explained by a linear model that aims to predict log2-transformed fold changes in the listed ChIP-seq read counts between myoblasts and myotubes. Two models are considered. The first, with performance indicated as gray bars, uses a site’s accessibility and MYOD binding status. The second, indicated as black bars, augments the first with accessibility and MYOD at linked sites. See Methods for more details.

**Supplemental Figure 6.**
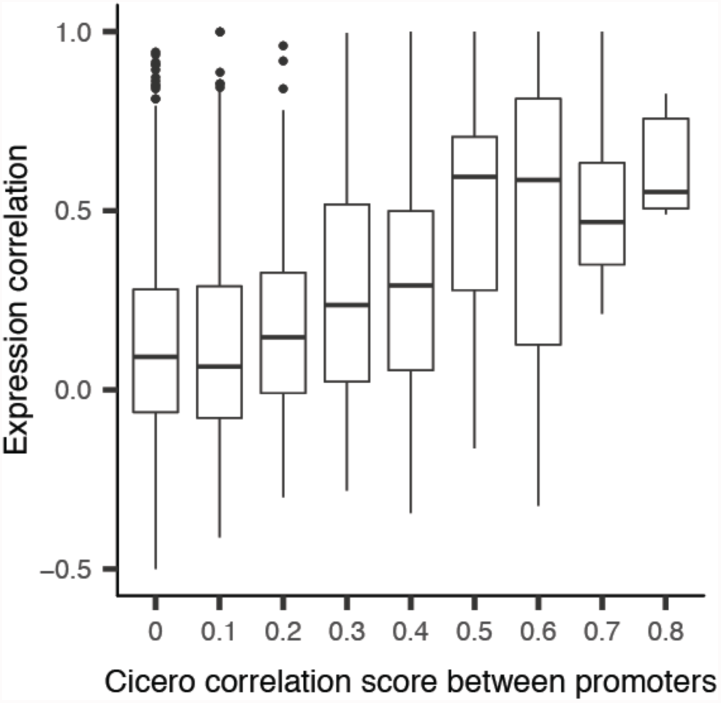
Correlation in expression among linked differentially expressed genes. Boxplots of the cell-wise correlation between gene expression among pairs of differentially expressed genes whose promoters have different Cicero co-accessibility scores.

